# A common contaminant shifts impacts of climate change on a plant-microbe mutualism: effects of temperature, CO_2_ and leachate from tire wear particles

**DOI:** 10.1101/2020.05.19.105098

**Authors:** Anna M. O’Brien, Tiago F. Lins, Yamin Yang, Megan E. Frederickson, David Sinton, Chelsea M. Rochman

**Author notes:** Corresponding author Anna M O’Brien: Anna M O’Brien Department of Ecology & Evolutionary Biology 25 Willcocks St Toronto, Ontario, Canada, M5S 3B2.

## Abstract

Anthropogenic stressors, such as climate change or chemical pollution, affect individual species and alter species interactions. Moreover, species interactions can modify effects of anthropogenic stressors on interacting species - a process which may vary amongst stressors or stressor combinations. Most ecotoxicological work focuses on single stressors on single species. Here, we test hypotheses about multiple stressors (climate change and tire wear particles) and interacting species, and whether species interactions modify responses. We use duckweed and its microbiome to model responses of plant-microbe interactions. Climate change is occurring globally, and with increasing urbanization, tire wear particles increasingly contaminate road runoff. Their leachate is associated with zinc, PAHs, plastic additives, and other toxic compounds. We crossed perpendicular gradients of temperature and CO_2_ in a well plate with factorial manipulation of leachate from tire wear particles and presence of duckweed microbiomes. We measured duckweed and microbial growth, duckweed greenness, and plant-microbe growth correlations. We found that tire leachate and warmer temperatures enhanced duckweed and microbial growth, but microbes diminished positive responses in duckweed, meaning microbiomes became costly for duckweed. These costs of microbiomes were less-than-additive with warming and leachate, and might be caused by leachate-disrupted endocrine signaling in duckweed. We observed reduced greenness at higher CO_2_ without tire leachate, suggesting a relative increase in plant nutrient demand, and possibly underlying positive plant-microbe growth correlations in these conditions, as microbes presumably increase nutrient availability. However, with tire leachate, growth correlations were never positive, and shifted negative at lower CO_2_, further suggesting leachate favors mutualism disruption. In summary, while individual stressors of global change can affect individual species, in ecology we know species interact; and in ecotoxicology, we know stressors interact. Our results demonstrate this complexity: multiple stressors can affect species interactions, and species interactions can alter effects of multiple stressors.

## Introduction

Global change can disrupt species interactions. Famously, rising temperatures cause corals to expel mutualistic symbionts (Hoegh-Guldberg, 1999), and eutrophic conditions cause cascading effects on lake food chains (Carpenter et al., 2001) and decouple fitnesses in nutrient exchange mutualisms (Shantz et al., 2016). With the increase of human influence, global change extends beyond CO_2_, temperature, and nutrients, as these factors are now matched or exceeded in rates of increase by synthetic contaminants (Bernhardt et al., 2017). Despite proportionally less attention in the ecological literature (Bernhardt et al., 2017), synthetic contaminants have similarly far-reaching impacts on species interactions and food webs. Upon exposure to ozone, the anti-herbivory benefits of hosting a fungus disappeared for plants (Ueno et al., 2016), and certain groups of synthetic contaminants shift rates and diversity of whole clades of parasites (Blanar et al., 2009). Contaminants can also have pervasive indirect effects via trophic cascades in aquatic ecosystems (Fleeger et al., 2003). A synthetic oestrogen in a lake caused a prey fish species to crash, indirectly reducing the top predator and increasing zooplankton biomass (Kidd et al., 2014). Importantly, synthetic contaminants should be considered in the suite of global change stressors.

Species interactions may also alter the individual species-level effect of stressors - including chemical contaminants. In addition to affecting a physiological response, they can alter the dosage and/or mechanism of exposure that individuals receive. From DDT to microplastics, pollutants can move to predators from prey via consumptive interactions (Hickey and Anderson, 1968; Nelms et al., 2018). Likewise, mutualistic rhizosphere microbes can cause higher concentrations of heavy metals in plant tissues than the plant would accumulate alone (Braud et al., 2009). Direct effects of contaminants on one species can also combine with the effects on species interactions, i.e. to increase or decrease the rates at which species interact, and therefore the rates of trophic transfer. For example, neonicotinoid pesticides reach and harm non-target insect predators through consumption of contaminated prey, which reduces predation pressure and leads to a population increases of tolerant prey (Douglas et al., 2015). Contaminants may also change physiology and behaviors, producing trait-mediated shifts in interactions (Saaristo et al., 2018), such as psychoactive pharmaceuticals that change predator avoidance behavior and therefore predation rates on fish (Martiny et al., 2013).

We have a vested interest in the outcomes of certain species interactions. Plant-microbiomes are broadly tied to human well-being through influences on ecosystem productivity, crop health, and even on our own microbiomes via vegetable consumption (Berg et al., 2014). Further, a subset of plant-microbe interactions supply the majority of terrestrial plant nitrogen and phosphorus (Smith et al., 2011; Fowler et al., 2013; Coskun et al., 2017). Plant-microbiome interactions are largely mutually beneficial (Avis et al., 2008; Dijkstra et al., 2013), and often ameliorate stressors (Porter et al., 2019), including temperature and drought (Compant et al., 2010; Kivlin et al., 2013), yet can alternatively exacerbate negative effects (David et al., 2018). While microbes often dilute contaminant effects on plants because they either promote growth (Rajkumar et al., 2012), or reduce uptake by metabolizing or bioadsorbing compounds (Chaudhry et al., 2005; Madhaiyan et al., 2007), some microbes instead enhance effects by increasing contaminant bioavailability (Braud et al., 2009). Effects of synthetic contaminants on plant-microbe mutualisms are poorly understood, and here, we use interactions between duckweed *Lemna minor* and its microbiome as a model. Duckweed has an extensive history in ecotoxicology, owing to its ability to adsorb or transform a wide variety of anthropogenic contaminants, from heavy metals (Mo et al., 1989), to nutrients (Zhao et al., 2014) and organic compounds (Gatidou et al., 2017; O’Brien et al., 2019). Duckweed has also proven to be a highly tractable experimental system due to its clonal reproduction, small size (a few mm, Landolt, 1975), short generation time (as few as 3 days, Liu et al., 2017), and host-microbiome interactions similar to those of other plants, in which microbiomes promote duckweed growth in benign and stressful conditions (O’Brien et al., 2019; O’Brien et al., 2020*b*,*a*).

We aim to quantify the effects of a single chemical stressor, tire wear particles, on duckweed-microbiome interactions. Recent estimates suggest a massive 1 million t/a of tire wear particles enter the environment in the USA, with increasing inputs in recent decades due to mounting vehicle traffic (Wagner et al., 2018). Tire wear particles are the main source of total suspended solids (Göbel et al., 2007), zinc (Councell et al., 2004), and polycyclic aromatic hydrocarbons (PAHs, Boonyatumanond et al., 2007) in urban runoff, and leachate from tires has been linked to acute lethality in coho salmon (Peter et al., 2018) and developmental abnormalities in fathead minnow (Kolomijeca et al., 2020), but has shown milder effects on other organisms (Marwood et al., 2011; Panko et al., 2013; Redondo-Hasselerharm et al., 2018). Since multiple stressors very often underlie “ecological surprises” (non-additive effects e.g. Darling and Côté, 2008; Crain et al., 2008; Jackson et al., 2016), and since the multiple facets of global change do not occur in isolation, we consider effects of tire wear particles across gradients of climate change. We evaluate these global change factors for effects at multiple levels, from single-stressor on single-species, to multi-stressor on interaction out-comes, and we specifically consider how shifts in variation within interaction outcomes could alter long-term responses. In mutualisms, fitness feedbacks (correlations between fitnesses of interacting species, i.e. Sachs et al., 2004), might shift with environmental conditions (Shantz et al., 2016), with positive fitness feedbacks enhancing mutualisims, and weak or negative feedbacks potentially leading to evolutionary disruptions (Weese et al., 2015).

The well-documented positive effect of CO_2_ on both microbial and plant growth (Treseder, 2004; Norby and Zak, 2011), is linked to increases in root exudates (Phillips et al., 2006), which appear to drive microbial growth responses that in turn enhance nitrogen turnover and feed back to plant growth (Phillips et al., 2011). Thus, we predict that elevated CO_2_ will enhance the main benefits of microbes to duckweed and investment by duckweed in microbes, as well as enhance positive plant-microbe fitness correlations. Likewise, the impacts of microbes on plant growth in response to increases in temperature are most often positive, even if the main effects of temperature are sometimes not (Compant et al., 2010; Kivlin et al., 2013), therefore we expect temperature could also enhance both positive effects of interactions and fitness correlations. Conversely, predicting effects of leachate from tire wear particles is less straightforward, as the two chemicals often implicated in leachate effects, zinc and PAHs, would be expected to cause contrasting responses. Elevated zinc levels negatively affect duckweed, and while its microbiome can reduce negative impacts in the short term (O’Brien et al., 2020*b*), long-term benefits of microbiomes may erode, as zinc may cause negative fitness feedbacks between duckweed and microbes (O’Brien et al., 2020*a*). In other systems, microbes often enhance uptake of metal contaminants under warming and CO_2_ (Rajkumar et al., 2013), suggesting that climate change may exacerbate both short- and long-term effects of zinc. Yet for PAHs, microbes may mitigate negative effects: PAHs generally have negative effects on plants, including duckweed (Becker et al., 2002; Zezulka et al., 2013), but may be rapidly degraded by microbes (Heitkamp and Cerniglia, 1987; Haritash and Kaushik, 2009). While zinc and PAHs are most often expected to drive biological effects, leachate from tires contains a complex mixture of compounds (Peter et al., 2018; Kolomijeca et al., 2020; Capolupo et al., 2020) and main effects on responses of organisms are highly varied (Panko et al., 2013; Peter et al., 2018), precluding clear predictions. However, several studies have found greater effects of leachate at higher temperatures (Marwood et al., 2011; Kolomijeca et al., 2020), so we might predict that warming would exacerbate leachate effects on duckweed, microbes, and their fitness correlations.

## Methods

### Biological materials

We collected *Lemna minor* and associated microbes from the University of Toronto’s field station, the Koffler Scientific Reserve (King City, Ontario, Canada), in the summer of 2017. We used one single frond (bleached to remove all source-site microbes) to start an isogenic line (or nearly so), which grew to high numbers (>1,000) in just a few months. *Lemna minor* is known to reproduce primarily via vegetative clonal budding of daughter fronds but it can also very occasionally undergo sexual reproduction (Ho, 2017). Although we never observed sexual reproduction in the lab, flowers are cryptic (Landolt, 1975). Even if sexual reproduction occurred in our cultivated line, given little segregating variation within duckweed from our source site (Ho, 2017), we expect our line is still essentially isogenic, and assume so from here forward. Plants in our isogenic line were cultured in growth chambers (ENCONAIR AC80, Winnipeg, Canada) under 16-hour 23 ΰC day and 8-hour 18 ΰC night cycles, in Krazčič’s media (Krazčič et al., 1995), in vented 500 mL mason jars. We refreshed media approximately twice per month, as the isogenic line was maintained at high density. When duckweeds were collected, we also isolated the microbes associated with them by pulverizing one clonal unit of duckweed (e.g. a mother and daughter frond), plating on yeast-mannitol agar media, and culturing at 29 ΰC for 5 days before storing at 4 ΰC until the experiment. This microbial culture represents the fraction of culturable microbes in both the external and internal microbiome (e.g. epiphytic and endophytic), and includes a subset of bacterial taxa that are representive of the field sampled bacterial microbiome (O’Brien et al., 2019; O’Brien et al., 2020*b*), but also may include other taxa, as fungi and diatoms are known to associate with duckweed (Rejmankova et al., 1986; Goldsborough, 1993), and may have persisted in lab cultures. Previous sequencing of the bacterial fraction of the microbial culture identified 17 unique members, largely from Gammaproteopacteria (but also Alphaproteobacteria, Bacilli, Flavobacteriia, Firmicutes, and Sphingobacteriia), with Aeromonadaceae (including *Aeromonas* spp.) and *Pseudomonas* spp. in relatively high abundance (O’Brien et al., 2019).

Three to four days before adding duckweeds to the experiment, we sterilized the external surfaces, as our cultures are vented to lab air and not gnotobiotic. We shook for 5 minutes in reverse osmosis water, submerged them in 1% NaOCl (diluted from Lavo Pro™, Montréal) for one minute in a biosafety cabinet, then rinsed with autoclaved water four times: the first rinse short and vigorous to remove most bleach, then three 10 minute submerged soaks. While this procedure does not remove all endophytic microbes inside tissues, we have found that it is successful at greatly reducing microbes (Figure S1, O’Brien et al., 2019), and that duckweed generally does not recover from longer or more concentrated bleach treatment (O’Brien, pers obs).

### Experimental Device

We used an experimental device for our multi-stressor and fully factorial experiment that was based on previously reported designs applying CO_2_ gradients over multi-well plates (Nguyen et al., 2018; Yang et al., 2020). Here, we included orthogonal temperature manipulation, added gas delivery tubes to allow internal lighting, improved the CO_2_ concentration control system, and humidified the air (see Figure S2, for a graphical representation).

In brief, CO_2_ from a gas tank (Praxair) was manually adjusted via regulator and needle valves in a hand-assembled gas mixing board. The control system maintained a constant concentration of CO_2_ in ppm, as described by Yang et al. (2020), and was comprised of an Arduino microprocessor connected to a solenoid valve, a 12V/5V relay module, and a CO_2_ sensor. The valve and relay were controlled by a PID algorithm in response to sensor output within the CO_2_-air mixer, maintaining a steady concentration of CO_2_ into the experiment by turning the flow on and off. The CO_2_ concentration in the CO_2_ rich stream was monitored continuously using the Arduino user interface. We humidified CO_2_-rich air and ambient air in separate hand-assembled bubble humidifiers, which forced gas into water-submerged air-stones. Finally, we pumped (Pawfly Adjustable Air Pump 4-LPM) humidified CO_2_-rich air and ambient air into opposite sides of each aerogel bar, developing a spatially linear gradient of CO_2_ concentration (Nguyen et al., 2018; Yang et al., 2020) from 1,000 to 400 ppm (across ranges from RCP8.5, USGCRP et al., 2017).

Since we aimed to quantify both independent and combined effects of climate change variables, we applied a thermal linear gradient orthogonal to the CO_2_ gradient. We used aluminum plates cut to fit 96-well plates, with aluminum tubes attached below the aluminum plate at both sides with thermal adhesive tape. We pumped (Esky EAP-03 2500L/H Submersible Water Pump) hot water through the tube under the plate on one side, and cold water on the other using flexible tubing from hot and cold tanks, heated (Anova Precision Cooker 4.0) to 35 ΰC and cooled (Active Aqua Chiller Refrigeration Unit, 1/10HP) to 7.2 ΰC. We monitored temperature daily throughout the experiment with thermocouples (Omega HHP806), and achieved a realistic global change temperature gradient based on the range of July stream water temperatures from rural to urban sites in the Greater Toronto Area (13-27 ΰC, Toronto Regional Conservation Authority, 2016). Temperature periodically fluctuated due to cycles in lab temperature that made chilling more and less effective, and also due to periodic leaks and tube-blockages in the system, which we corrected as they appeared.

We supplied light to the experiment by connecting experimental well plates to the aerogel gas gradient via delivery tubes with interlaced lighting. Delivery tubes consisted of two PCR 96-well plates, each with the tube ends removed with a hot wire foam cutter, with the second plate inverted and the open tube ends of the second pressed inside the open tube ends of the first. LED striplights (LEDMO 6000K 2835 SMD LED) were placed between each row of PCR plate wells (twice per row and including outside edges, so that each experimental row received light from both sides). The “top” side of one PCR was placed on top of the experimental 96-well plate opening, and the “top” side of the other against the aerogel. LED lights were set to their lowest brightness setting via a LED controller (ER CHEN) and emitted light parallel to the experimental plate liquid surface (indirect). All layers were pressed together with 3-1/2’ steel screws and nuts, and possible leakage of gas exiting directly from the well plate (rather than exiting out the aerogel as intended) was slowed by wrapping the device with parafilm.

### Tire wear particle leachate

We used a Michelin energy saver tire (a/s, all season, sidewall markings 205/60R16 91Vtire), and sliced strips from the tread portion, which we hand cut to small cubes (≈0.5 cm^2^). We ground cubes in a Cuisinart Supreme Grind Automatic Burr Mill (DBM-8C) with plate grinders, freezing the tire sample in liquid nitrogen before each grinding, and passing all tire particles through each setting (from most coarse to most fine). We then passed resulting particles through a burr mill with cone grinders (10903-913US, BODUM) three times, on the finest setting and at room temperature, which ripped tire pieces to provide a surface texture similar to road wear.

We characterized the size distribution of our lab-created tire wear particles by placing a 50 mg subsample in surfactant (10% w/v Alcojet, Alconox, Inc) on a glass slide. Particles were largely aggregated without surfactant, and some even with surfactant (Figure S3). We imaged all portions of this slide with a Leica (M205 A) microscope with camera (DFC425 C), at gain 1, gamma 0.57, whites blown out (“cut” to 25 or 26), and an automatic exposure time (ranged from 38.5 to 46.8 ms), with an added scale bar, using Leica Application Suite (version 3.8.0) software. We used ImageJ to analyze the number and size distribution of particles, with brightness thresholding (set to 195) and “Analyze Particles” to measure maximum caliper and top (facing camera) surface area. Our particle size distribution is coarsely similar to many measured size distributions for tire particles generated by road wear (Kreider et al., 2010; Wagner et al., 2018). We estimated 11.7 particles per milligram. The maximum caliper of particles ranged from 1.7 *μ*m to 1.7 mm, and facing surface area from 0.002 *μ*m^2^ to 0.9 mm^2^ (Figure S4), with particles generally in highly complex shapes (Figure S3).

Tire particles do enter waterways (e.g. Grbić et al., 2020), yet highly concentrated leaching near the roadside with dilution in recipient streams is expected to be the primary source of tire leachate contaminants, as the bulk of tire particles seem to remain near the road-side (Wagner et al., 2018). We sought to mimic this process. We leached tire particles immediately prior to use at a concentration of 20 g/L. Leaching took place in amber bottles wrapped in foil in autoclaved reverse osmosis water, with paired bottles for with (leachate) and without (negative control) tire particles. We set bottles on a shaker at 20 rpm for 10 days at ambient temperature. Leachate and negative controls were filter-sterilized (auto-claving would alter chemistry) with water-wettable polytetrafluoroethylene filters of 0.2*μ*m maximum pore width (Acrodisc® syringe filters, Pall Corporation, NY, USA) which in turn were sterilized by passing 1% NaOCl in sterile reverse osmosis water through the filter and letting it sit for 5 minutes, followed by triple rinsing with sterile reverse osmosis water. We expect that filtering removed the majority of tire particulate, limiting any observed effects to the leached chemicals. Still, while we did not detect tire particles smaller than 0.2*μ*m in diameter in our image analysis (minimum 1.7*μ*m, Figure S4), any that existed would have passed through this filter. We then split leachate into two solutions, one undiluted and one diluted by 50% with the negative control. To each experimental well, we added 100 *μ*L of a leachate treatment (negative control, 50% diluted, or full strength) depending on the design (see below), and we also added 100 *μ*L of double strength Krazčič’s growth media. Each well then holds 200 *μ*L of 1 × strength Krazčič’s media with a concentration of leachate that is 0, 0.25 ×, or 0.5 × the original leachate concentration. Our 0.5 × leachate treatment is meant to replicate the max reasonable dosage that a pond near a highway might receive, 10 g/L of leaching tire wear particles, and our 0.25 × treatment a dose that a less-travelled road or further pond might receive, 5 g/L of leaching tire wear particles (see ranges in Wagner et al., 2018). Leaching rates from tire wear particles may differ across leaching concentrations, but we expect these effects to be small (Rhodes et al., 2012), and so refer to our treatments as 5 and 10 g/L leachate throughout.

### Experimental set up

We experimentally exposed duckweed in well plates to climate-change gradients, crossed with tire leachate and microbial treatments. We used our device to generate perpendicular temperature and CO_2_ gradients over plates, so that each well in a plate is a unique combination of temperature and CO_2_ ppm. Within plates, each well received one sterilized duckweed clonal unit (e.g. one mother-daughter frond pair), and we alternated microbial re-inoculation across columns of wells, so that both re-inoculated and uninoculated treatments spanned the temperature gradient.

To generate inocula, we placed a swab from the cultured agar plate (see above) into 2 mL of autoclaved liquid yeast mannitol media in a glass vial (previously cleaned and sterilized in a muffle furnace) and cultured at 30ΰC for approximately 24 hours at 200 rpm in a shaking incubator (VWR, Radnor, PA, USA), together with an identical vial of yeast-mannitol media with no swab added as sham inoculum. We then diluted so that 10 *μ*L of inocula would bring a well to approximately 5,000 cells per *μ*L based on an estimate of cell density from optical density (as described in O’Brien et al., 2020*b*). We diluted sham inocula by the same amount, added 10 *μ*L of inocula or sham to each well, and sealed plates with BreatheEasier (Millipore-Sigma, Diversified Biotech, Dedham, MA, USA) membranes. We crossed this design with three levels of tire leachate (96 unique temperature and CO_2_ conditions × 3 = 288 treatments) and treated an entire plate with a particular level (0 g/L = none, 5 g/L, or 10 g/L, see above) to prevent cross-contamination between treatments (Birch et al., 2019). We expect that many components of tire leachate could be volatile (U.S. EPA CDC/ATSDR, 2019), yet we require gas exchange for CO_2_ treatments and living organisms. Plates in devices were connected to CO_2_, air, hot water, and cold water in parallel for 7 days. This three-plate setup constituted one replicate of the experiment, and we repeated the setup three times (i.e. 3 blocks, for 9 total plates and 864 total experimental units), where plates with different tire particle leachate treatments were randomly assigned to the three parallel devices within each replicate (see Figure S2).

All experiment setup including hand sterilization, plate filling, and microbial manipulation was conducted in a biosafety cabinet (ESCO Micro Pte. Ltd., Labculture®, Singapore) and glassware was used where possible, with standard cleaning followed by a muffle furnace treatment at 450ΰC for 7 hours (ThermoFisher Scientific, F30428C-80). We used glass-coated 96-well plates (Thermo Scientific, 60180-P304) that were bleach sterilized (5 minutes in 1% NaOCl, followed by three rinses of autoclaved reverse-osmosis water) and cleaned of contaminants (three rinses of acetone followed by one rinse of hexane left to evaporate (both Fisher Chemical HPLC grade, ≥99.5% and ≥98.5%, respectively). When glassware was not possible, plastic labware was autoclaved or purchased pre-sterilized.

### Data collection

At the end of the experiment, we disconnected plates from the device and recorded duckweed growth and traits with image analysis, and microbial growth with optical density.

We photographed plates using a custom camera rig for a Nikon D3200 (with AF-S DX NIKKOR 18-55mm f/3.5-5.6G VR lens, Minato, Tokyo, Japan) and a standard backlit lighting regime (created with Yongnuo YN-300 light, Shenzhen, China). We analyzed images in ImageJ, using color threshold settings to select only “live‘” duckweed fronds. Thresholds were set subjectively by the image scorer (blind to plate conditions) to include duckweeds having any green hue, but exclude algae. The same thresholds for minimum pixels and hue were applied across all plates in a round, but individual images differed slightly, so small adjustments to brightness and saturation cutoffs were necessary. We then measured pixel area and greenness (ratio of green brightness to total brightness in pixels across RGB channels) of all duckweed fronds in a well (with “Analyze Particles,” see examples in Figure S5). Greenness is associated with leaf nitrogen (Thind et al., 2012), including greenness in digital images (Rorie et al., 2011). This is likely due to the relationship between chlorophyll and nitrogen content (Schepers et al., 1992; Ma et al., 1996), which may be reduced by increasing CO_2_ (Ellsworth et al., 2004), as plants become nitrogen limited. We used a custom R script to sum (area) or average (greenness) measures for duckweed fronds in the same well.

Optical density was measured on a 70 *μ*L sample of liquid from each well at the end of the experiment after imaging. We recorded optical density at 600 nm (BioTek Synergy HT plate reader, Gen5 1.10 software, Winooski, VT, USA). Plates could not be measured simultaneously, and were incubated at 4ΰC 0.5-2 hours between imaging and optical density measures. Each reading had the optical density of the background (plate and reverse osmosis water) subtracted. The minimum optical density that was greater than 0 was taken to be the threshold reading, and all values lower than 0 were set to this threshold. We expect optical density to be correlated with the live colony forming units of microbes in solution at the end of the experiment, as verified in O’Brien et al. (2020*b*).

### Data analysis

We analyzed data in R (R Core Team, 2019) using linear models and with package MCMCglmm (Hadfield, 2010). We fit models from more complex (all linear interactions of all treatment parameters) to less complex in reverse stepwise regression. At each step, we removed the most complex (highest order interaction) parameter, unless more than one most complex parameter was non-significant, in which case we removed the term with the highest pMCMC (the Bayesian equivalent of the p-value, Hadfield, 2010). We repeated the process until no terms were non-significant (unless they were components of significant higher-order interactions, in which case they were retained), or the simpler model fit worse, and we call the resulting model the “best” model. We evaluated significance with pMCMC and model fit with the Bayesian equivalent of AIC, or deviance information criteria (Spiegelhalter et al., 2002). We report highest posterior density intervals (HPDIs) as the Bayesian equivalent of confidence intervals and use 95% for pMCMC <0.05 and 90% for effects with marginal pMCMC values (<0.1). All models included the random effect of round (1, 2, or 3), ran for 100,000 iterations thinned by 50 and with 500 iterations of burnin. We refit best models with increased iterations (1,000,000), for reporting estimated parameter values. We applied our best-model procedure to the response variables of duckweed growth in pixel area, the log of optical density (data from inoculated wells only; we fit log of optical density as a function of inoculation as a separate model, see Figure S1), and greenness.

For better behavior of model fitting functions, we modified CO_2_ concentrations to the same order of magnitude as temperature by dividing by 100 (ppm/100 or, parts per 10,000). We interpolated average measured temperatures over the course of the experiment for each well (due to slight variations, see above), and these differed somewhat in range across treatments and tire particle leachate treatments (wider in control, 13.8-29.3ΰC vs 19.0-26.5ΰC all others), but means were similar (21.7, 22.2ΰC for 0 g/L and all others, respectively).We do not anticipate that this affected the estimated main effect of leachate, given similar mean temperatures, nor do we expect that the wider range of temperatures in plates without leachate increased power to detect temperature effects in this vs other tire treatments, given that temperature effects are not strongest at 0 g/L (see Results). Finally, some wells partially dried due to loose seals with gas supply in the device. In these wells, duckweed generally died due to sticking to well walls as liquid levels dropped, rather than due to treatment exposure, and optical density-based estimates of microbial growth will be inaccurate. These wells were difficult to identify with certainty in some cases, so we excluded all wells in which duckweed had fully died (104 of 864 wells).

If microbes and duckweed independently respond to the same treatments, this could cause positive fitness correlations between the two without fitness feedbacks. However, we can use the residual fitness variation and covariation after accounting for the effects of treatments on duckweed and total microbial growth, to ask how selection might act on duckweed or microbe genotypes that increase the fitness of their partner. We further explored the residuals of duckweed and microbial growth from a fully fitted model with all terms, including nonsignificant treatment terms (in case weak trends were missed in best models), and using the data from only inoculated wells in which duckweed survived (n=372). We modeled residuals of duckweed pixel area increase as the response, and explanatory variables were residual log optical density and the interaction between residual log optical density and each treatment or combination of treatments. As above, we performed reverse stepwise regression to select the best model.

## Results

Duckweed was affected by all anthropogenic stressors in either growth, traits or both. We also observed effects from interaction with the microbiome, and altered interactions in the presence of anthropogenic stressors, i.e., non-additive effects between the microbiome and anthropogenic stressors.

For duckweed growth, the best model found that microbiome effects were conditional (main effect pMCMC > 0.1) on both tire leachate and temperature (interaction terms pM-CMC < 0.05, Table 1, Figure 1), such that increasing tire leachate and temperature separately increased costs of microbiomes to duckweed. At both lower temperature and without any tire leachate, inoculated and uninoculated plants grew equally little (Figure 1b, see overlapping HPDI at lowest temperatures). Without any tire leachate, uninoculated plants responded positively to warming temperature (pMCMC < 0.05), but plants inoculated with microbiomes did not (pMCMC < 0.05), producing negative effects of microbiomes at warmer temperatures (Figure 1b). With 10 g/L tire particle leachate, uninoculated duckweed grew about three times as much as inoculated plants, on average across other treatments (means 2694 and 876 pixels, SE *±* 225 and 143 pixels, both respectively, Figure 1a, Tire ×Microbe negative with pMCMC < 0.05, Table 1), also producing costs of microbiomes. However, costs of microbiomes at higher temperatures and with tire leachate were less than additive, with marginal significance (Temperature ×Tire ×Microbes pMCMC < 0.1), meaning that plants without microbiomes did not grow much more than plants with microbiomes when temperatures were warmest and the tire leachate treatment was 10 g/L of particles (Figure 1c). The best fit model did not include an effect of CO_2_ on duckweed growth.

**Table 1:** Best fit models between treatments and response variables: duckweed growth, microbial growth (optical density), and duckweed frond greenness (proportion). CO_2_ was fit with ppm / 100 (or, parts per 10,000, ranging from 4-10), see Methods. “–” indicates this term is not in the best model for the response variable, “*” indicates pMCMC < 0.05 and “.” indicates pMCMC < 0.1

| parameter | duckweed growth | log optical density | greenness |
| --- | --- | --- | --- |
| Intercept | -2280 | -8.38 * | 0.400 |
| Microbes | 2410 . | NA | — |
| Temperature | 187 * | 0.151 * | 0.002 * |
| CO<sub>2</sub> | — | — | -0.003 * |
| Tire | 0.821 | 0.100* | -0.0007 |
| CO<sub>2</sub> × Tire | — | — | 0.0001 . |
| Microbes × Tire | -608 * | NA | — |
| Temperature × Tire | 3.16 | — | — |
| Temperature × Microbes | -152 * | NA | — |
| Temperature × Tire × Microbes | 24.4 . | NA | — |

**Figure 1:**
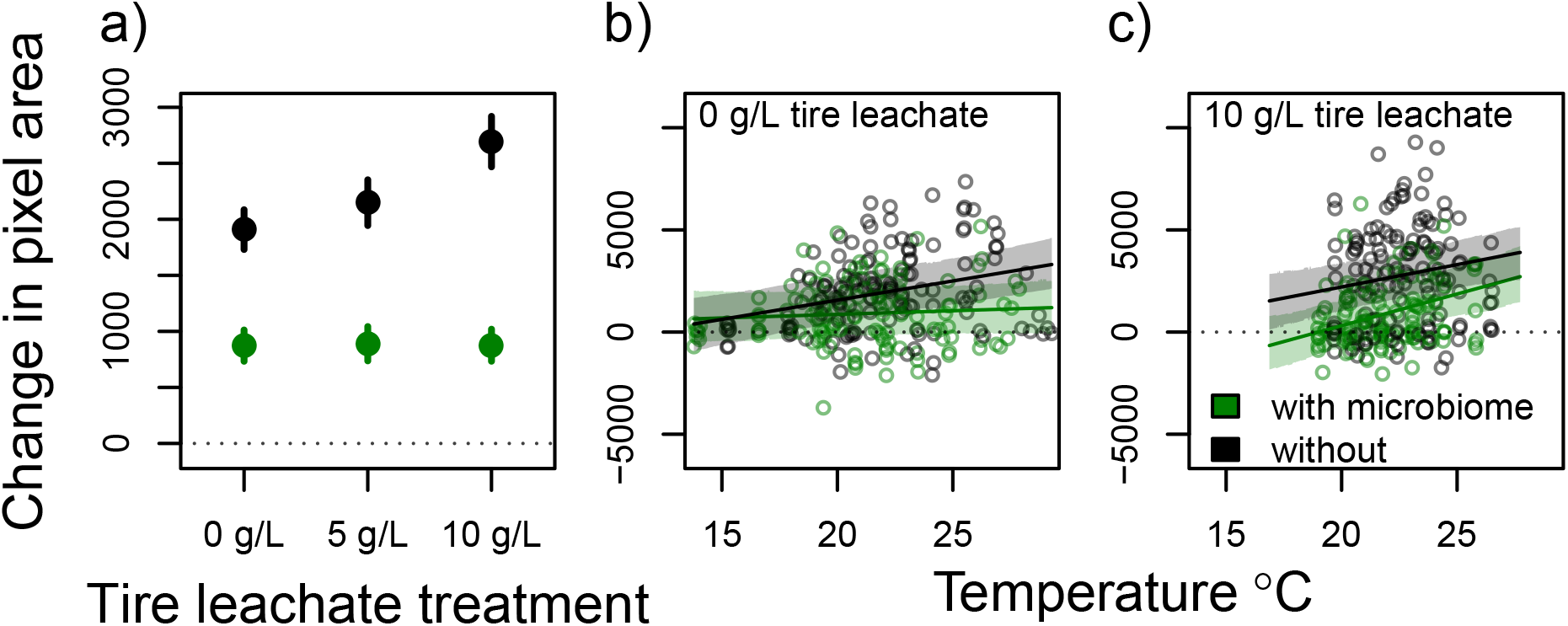
Duckweed growth when re-inoculated (green) and when not re-inoculated (black) with their microbiome across experimental treatments with significant effects. a) Growth means (points) across different levels of tire particle leachate treatments (x-axis), with one standard error of the mean (bars). b) & c) Duckweed growth across temperature (ΰC, x-axis), at 0 g/L (b) and 10 g/L (c) tire particle leachate treatments. Points are observed growth data with interpolated average temperature values during the experiment. Shaded backgrounds indicate 90% HPDIs for the predicted mean (lines) from the best-fit model. Data and fitted model predictions for concentrations of 5 g/L leachate treatments are not shown, but are intermediate. The temperature range extends further in (b) due to temperature anomalies in one round of the experiment.

Total microbial growth (putative average “fitness” across the microbiome community), was also affected by temperature (pMCMC < 0.05) and tire leachate (pMCMC < 0.05), but not CO_2_ treatments (not included in the best model). Like duckweed, microbes grew more at higher temperatures (mean predicted optical densities 0.003 and 0.025 at coldest and highest applied temperatures, respectively) and at higher tire particle leachate concentrations (mean predicted optical densities of 0.005 at 0 g/L and 0.015 at 10 g/L, non-overlapping 95% HPDI, Figure 2). There was some contaminant microbial growth in uninoculated wells, but there was less microbial growth, on average, in uninoculated wells (optical density means 0.0065 and 0.0090, with SE range 0.0061-0.0070 and 0.0083-0.0097 in uninoculated and inoculated wells, respectively, pMCMC < 0.05, Figure S1).

**Figure 2:**
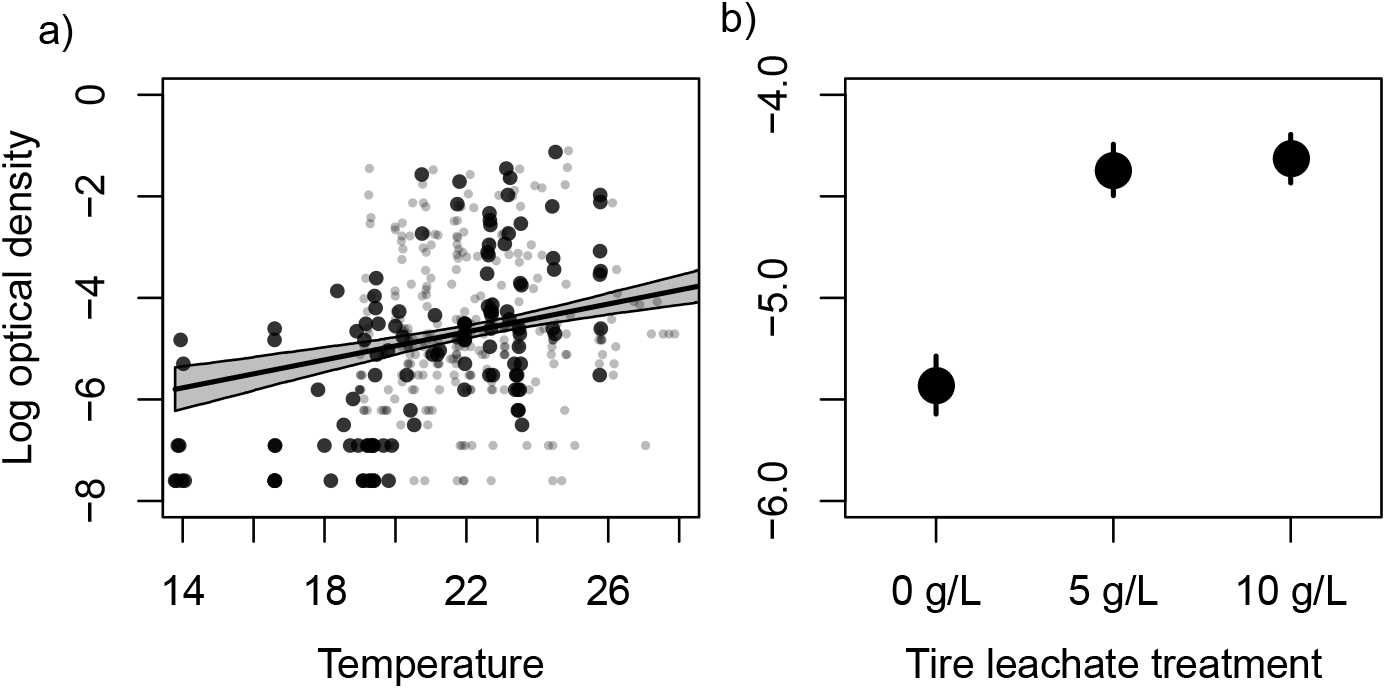
Response of total microbial growth to treatments in the best model. Total microbial growth (log of the optical density) is putative average “fitness” across microbiome component species. a) Points are observed optical densities and interpolated average temperatures (grey and smaller = 10 g/L or 0 g/L, black and larger = 5 g/L tire particle leachate treatments) for each well. Grey background indicates 95% HPDI for the predicted mean (line) from the best-fit model at 5 g/L leachate treatments. b) Means of logged observed optical densities (bars are standard errors) across 0 g/L, 5 g/L, and 10 g/L tire particle leachate treatments.

The best model for duckweed frond greenness found that CO_2_, temperature and leachate from tires all affected outcomes. Temperature increased greenness in wells by about 0.04 from coldest to warmest (from model-predicted mean of 0.408 to 0.440, Figure 3a, Table 1), similar to growth effects. In contrast, increasing CO_2_ from the lowest to highest applied level decreased greenness proportion by about 0.02 (from predicted mean of Figure 3b). Interestingly, with tire particle leachate, greenness was less reduced for higher CO_2_ levels, with only half as much change across the CO_2_ range, 0.01 (for 10 g/L, though still non-overlapping 95% HPDIs between lowest and highest CO_2_).

**Figure 3:**
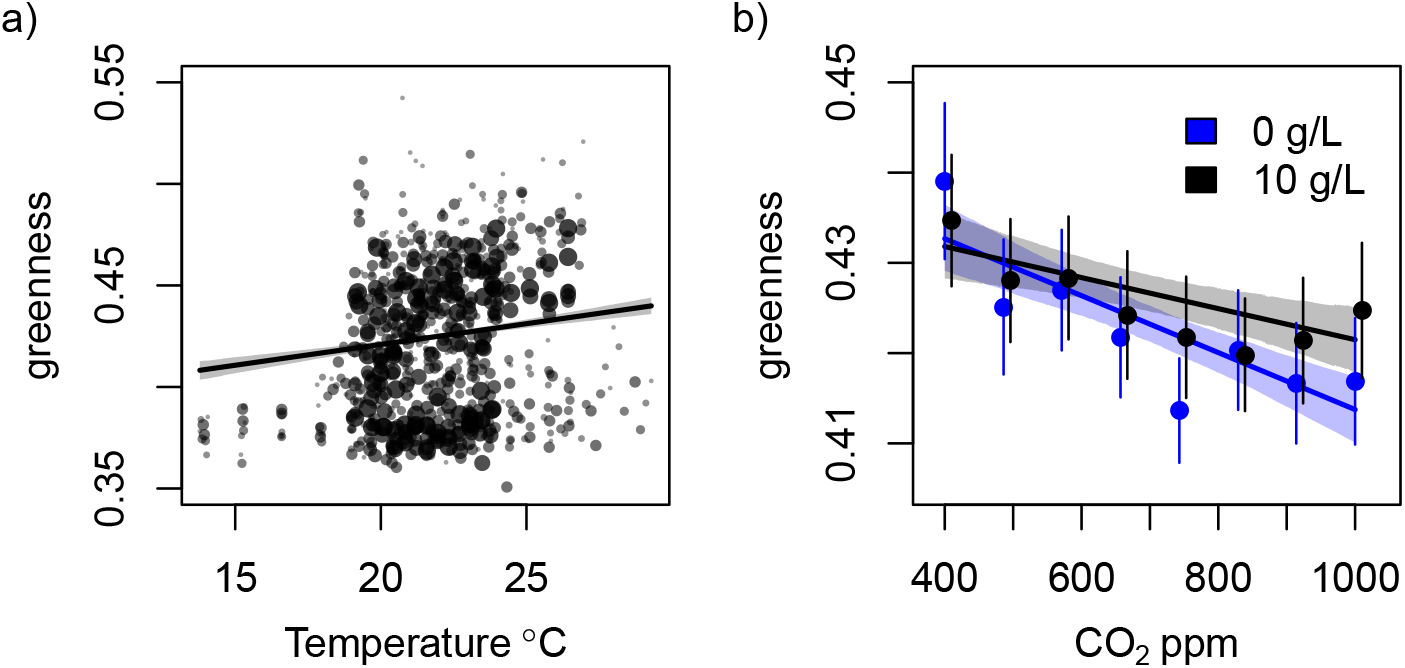
Response of greenness of duckweed fronds to treatments. a) Points are observed greenness across interpolated average temperatures. Background shading indicates 95% HPDI for the predicted mean (line) from the best-fit model for the parameters tested, at 700 ppm CO_2_ and 5 g/L tire particle leachate treatments. Points from treatments further from these values are smaller and fainter. b) Greenness in CO_2_ and tire particle leachate treatments (blue, 0 g/L and black, 10 g/L) averaged across temperature (bars are one standard error), offset slightly along the x-axis for visibility. Model predictions (lines) and 95 % HPDI (shading) are for average temperature. Microbiome treatment did not affect greenness, so observations in both panels are included as points (a) or averages (b) without regard to microbe treatment.

Raw fitness correlation was essentially neutral (*ρ*=-0.016), despite similar responses of duckweed and microbes to treatments, which might have inflated correlation. Indeed, residual fitness correlations accounting for main treatment responses were largely negative, and the best fit model suggested significant influence of experimental treatments on the sign and strength of fitness correlations. There was a marginally significant negative correlation between residual duckweed and total microbial growth. This was shifted positive by CO_2_ (marginally, pMCMC < 0.1) and temperature (n.s.), but further decreased in higher tire particle leachate concentration treatments (marginally, pMCMC < 0.1). While effects were all marginally or not significant, all simpler models (factorial combination of remaining terms, and intercept only model) fit worse when evaluated by DIC. We visualized these marginally significant effects by splitting the data into two scenarios for weaker and stronger climate change (temperature and CO_2_ above or below the average level we applied, 21.7ΰC and 700 ppm, respectively) crossed with no, medium, and higher prevalence of tire wear particles. With weaker climate change conditions (13.7-21.7ΰC and ambient-700 ppm CO_2_), residual fitness correlations were marginally negative without leachate from tire particles, but strongly negative (significant, HPDIs for duckweed growth do not overlap for extremes of residual microbial growth) with 10 g/L leachate (Figure 4a-c). Under stronger climate change conditions (21.7-29.3ΰC and 700-1000 ppm CO_2_), fitness correlations were marginally positive without leachate, but shifted to neutral with 10 g/L leachate treatments (Figure 4d-f).

**Figure 4:**
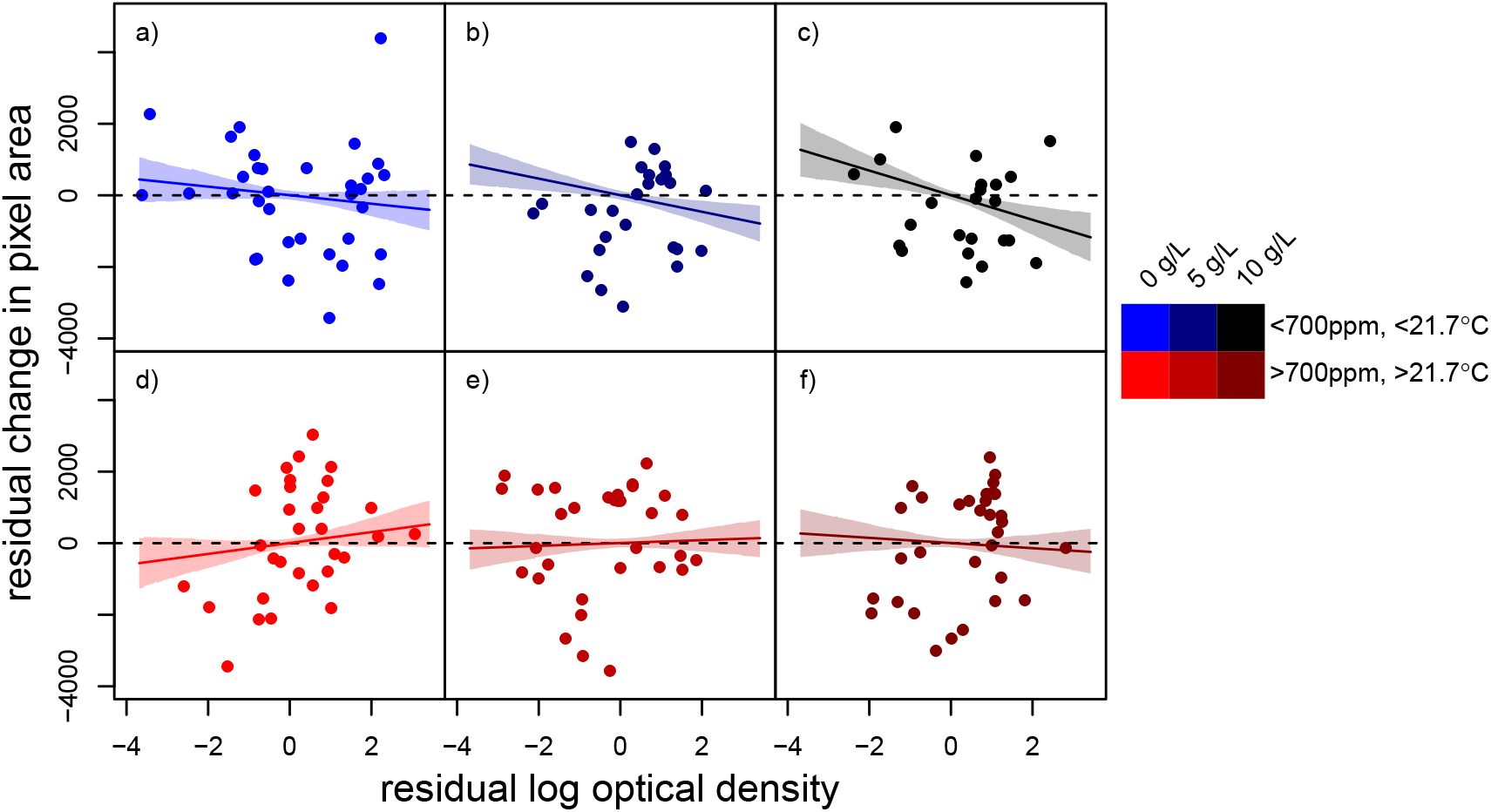
Residual growth (fitness proxy) for duckweeds and microbes for subsets of data falling into scenarios of weaker (a - c) or stronger global change (d - f) with 10 g/L (c and f), 5 g/L (b and e) and without (a and d, 0 g/L) contamination with leachate from tire wear particles. Stronger global change is defined as CO_2_ and temperature both above average treatment means, 21.7ΰC and 700 ppm, and shown in reds. Weaker global change is defined as CO_2_ and temperature below these means, and shown in blues. Tire particle leachate treatment is indicated by color darkness, with brightest colors indicating no leachate, and darkest indicating 10 g/L. Points are data observations from these treatment categories, with shaded background indicating the 90% HPDI for the predicted mean response (lines) of the best fit model at the mean of the treatment ranges within the category. Strong warming with low CO_2_ and vice versa represent less likely global change scenarios and data and model predictions are not depicted here.

## Discussion

### Climate change and tire wear particle leachate have non-additive effects on duckweed-microbiome interactions

Our observations show the importance of considering synthetic contaminants in global change science. Synthetic contaminant concentrations in nature, including pesticides, plastic additives, and trace metals, are prevalent and increasing - and in step with other global change parameters (Bernhardt et al., 2017). As shown here, synthetic contaminants can have effects that can both percolate through species interactions (Fleeger et al., 2003) and shift across climate change backdrops (Yang et al., 2020). We aimed to characterize the simultaneous and individual effects of climate change and a model synthetic contaminant (leachate from tire wear particles) on duckweed and its microbiome. Leachate from tire wear particles altered duckweed growth, microbial growth, and duckweed-microbiome interactions, but effects varied with different climate change parameters. Both duckweed and microbes grew better under conditions increasingly resembling urban and future scenarios, i.e. warmer and more concentrated leachate from tire particles, but effects were not linearly additive for duckweed when microbes were present. We have previously found that duckweed microbiomes generally increase duckweed growth in both benign and a variety of stressful conditions (O’Brien et al., 2019; O’Brien et al., 2020*b*,*a*). Despite our strong prior that the relationship between duckweed and its microbiome would be mutualistic and would increase the benefits of CO_2_, microbes largely reduced the positive effects of temperature and leachate (but less so for combined warm temperatures and concentrated leachate, Table 1 and Figure 1), and there was no main effect of CO_2_ on growth of either duckweed or microbes. In other words, global change scenarios increased the costs of microbes as a whole relative to the benefits, shifting the interaction from essentially neutral at current, non-urban conditions (no leachate, low temperature, Figure 1b), to costly in future scenarios.

### Growth correlations, greenness and longer-term impacts on duckweed-microbiome interactions

The microbiome-free state may be irrelevant in nature: as no genotype of duckweed is microbe-free in the field, it cannot be favored by selection. Instead, variation in the quantity of microbes supported by hosts is a more meaningful metric (Partida-Martinez and Heil, 2011). In sections of the experiment representing only weak global change (Figure 4a), duckweed and microbe residual growth (variation after accounting for responses to temperature and tire particle leachate) were not positively correlated (Figure 4a). Stronger climate change manipulations (temperature and CO_2_ increases only) increased positive growth correlations (Figure 4d), but leachate from tire wear particles shifted correlations negative (Figure 4c,f). Growth correlations may indicate fitness conflicts and alignment in mutualisms (negative and positive correlations, respectively), and phenotypic correlations can be a good proxy for genetic correlations (Waitt and Levin, 1998; Steppan et al., 2002). However, this proxy is not always reliable (Stinchcombe et al., 2002) and may somewhat depend on genetic variation contributing substantially to phenotypic variation, and here we have only one genotype. While even clonal duckweed can accumulate mutations on which selection could act (Ho, 2017), our experimental duckweed are an unknown and relatively small number of clonal generations apart. Still, the relationship between growth of plants and the growth of microbes may be mechanistically the same regardless of whether variation is generated by stochastic or genetic effects. If so, over longer time, climate change could select for enhanced duckweed-microbiome mutualisms in uncontaminated sites, but for disrupted mutualisms in sites that receive road runoff.

The effects of treatments on plant greenness may help explain patterns, and provide a common mechanism that could link stochastic and genetic fitness correlations. Greenness is positively linked to leaf nitrogen and chlorophyll content (Rorie et al., 2011; Thind et al., 2012), and is often decreased when plants become nitrogen limited at elevated CO_2_ (Ellsworth et al., 2004). Therefore, reduced greenness may be a signal that plants are more nitrogen limited than carbon limited, and that microbial provisioning of nitrogen would have more fitness benefits and microbial use of carbon fewer fitness costs. Here, duckweed greenness was low with high CO_2_ when there was also no tire leachate pollution, and indeed, these same conditions (high CO_2_ and no tire leachate) were the only conditions in which microbe and duckweed growth were positively linked (Figures 3b & 4c, Table 2), suggesting microbes alleviate the nitrogen limitation of duckweed at elevated CO_2_. Indeed, relative strengths of carbon and nitrogen limitation appear to drive results across many different experimental manipulations of CO_2_ (Treseder, 2004; Phillips et al., 2011).

**Table 2:**
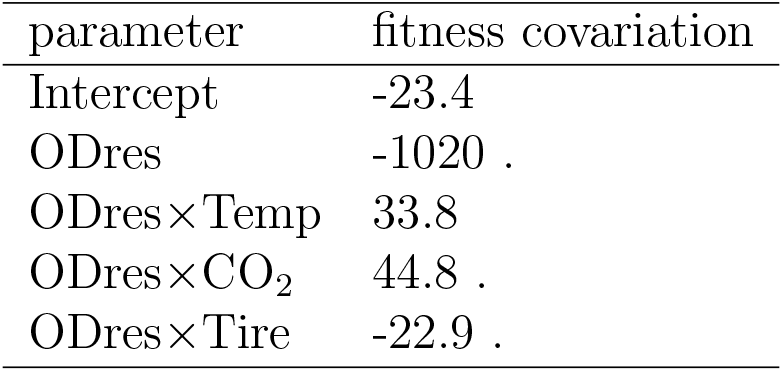
Best fit models for the slope between fitness residuals across treatments. “ODres” is short for the residuals of the log of optical density, and “.” indicates pMCMC < 0.1.

| parameter | fitness covariation |
| --- | --- |
| Intercept | -23.4 |
| ODres | -1020 . |
| ODres×Temp | 33.8 |
| ODres×CO<sub>2</sub> | 44.8 . |
| ODres×Tire | -22.9 . |

### Possible mechanisms of tire wear particle leachate effects

Another important question to consider is what components of the tire leachate might drive the effects we observed. Leachate of our exact tire is known to contain both zinc and PAHs (Kolomijeca et al., 2020), as does the leachate of many tires (Wagner et al., 2018). Zinc reduces duckweed growth at ambient climate (O’Brien et al., 2020*a*,*b*), so increased growth at higher leachate concentrations indicates that other components of tire leachate contribute to effects, masking or altering negative effects of zinc. One PAH is known to affect duckweed growth (phenanthrene, Becker et al., 2002), and a number of others inhibit growth and induce chlorosis in a close relative (anthracene, phenanthrene, benzo[a]pyrene, fluoranthene, pyrene and naphthalene, Huang et al., 1993; Ren et al., 1994). The tire we used contains all the above PAHs, and all but fluoranthene have been detected in its leachate (Kolomijeca et al., 2020). Surprisingly, we observed only positive effects on duckweed: increased growth and greenness at higher leachate concentrations (Figures 1a & 3b). Effects of PAHs may also depend on photodegradation pathways (Huang et al., 1993; Ren et al., 1994), and, interestingly, some effects of leachate from tires may likewise depend on light (Wik and Dave, 2006). Alternatively, very low doses of phenanthrene actually stimulated growth in duckweed (Becker et al., 2002), and increased growth at low doses due to compensatory mechanisms, a common biological response to toxins (Calabrese, 2008). Our growth responses could suggest low doses of PAHs in leachate, but the growth continues increasing from 5 to 10 g/L. More likely, PAHs could be disrupting hormone signaling in duckweed. Phenanthrene inhibits ethylene responses in *Arabidopsis thaliana* (Weisman et al., 2010), and as ethylene may promote or inhibit growth depending on the dose (Pierik et al., 2006), this could explain why effects vary across dose and plant species, beyond results here. In fact, plant endocrine disruption appears to be common in a number of classes of ubiquitous contaminants (Couée et al., 2013), and may be the mode of action here even if PAHs are not the component of leachate causing the observed effects.

Indeed, tires contain a great many other biologically active compounds (U.S. EPA CDC/ATSDR, 2019), many of which are known to leach into water (Zahn et al., 2019; Capolupo et al., 2020). Of those that are known to leach, mixtures of 1,3-diphenylguanidine and hexa(methoxymethyl)melamine are associated with toxic effects on coho salmon (Peter et al., 2018), and mixtures of benzothiazole, 2(3H)-benzothiazolone, phthalimide, phthalide, bisphenol-A, and n-cyclohexylformamide may underlie toxicity in algae and mussels (Capolupo et al., 2020). Much less is known about likely effects of these on duckweed. General toxicity of the transformation products of 1,3-diphenylguanidine is expected across organisms (Sieira et al., 2020), and the same is true for benzothiazole, with documented toxicity across a number of other species, and potential similar mode of action as PAHs, due to activation of aryl hydrocarbon receptors (Liao et al., 2018). Bisphenol-A toxicity to duckweed and other *Lemna* is known (Mihaich et al., 2009; Fekete-Kertész et al., 2015), and like PAHs may increase growth at low concentrations (Mihaich et al., 2009). Lastly, while hexa(methoxymethyl)melamine is expected to have low toxicity to aquatic organisms, (U.S. EPA, 2004) it has been associated with negative effects on *Daphnia* (de Hoogh et al., 2006). While the effects of these other compounds are largely expected to be negative, not enough is known to rule them out as sources of positive effects here. Future work should undertake non-targeted chemical analysis to ask whether there are unexpected chemicals in the leachate. Moreover, fractionating leachate could be included to determine which chemicals or suites of chemicals drive leachate effects on plants and microbes.

Regardless of the source of the growth promotion, microbes remove it, providing a clear example of a species interaction altering a contaminant effect. If duckweed responses to PAHs in tire leachate caused the increased growth, microbes may have removed effects by rapid mineralization. While we do not expect that the source of our biological materials has much history of contaminant exposure (field site in a natural area), even freshwater microbes from pristine sites may rapidly mineralize PAHs at appreciable rates (Heitkamp and Cerniglia, 1987; Haritash and Kaushik, 2009). PAH degraders include many *Pseudomonas* strains (Haritash and Kaushik, 2009), and several *Pseudomonas* strains have also been previously identified in this microbiome (O’Brien et al., 2019). This mechanism could also account for negative growth correlations induced by tire leachate: as microbes mineralize PAHs, they may increase in abundance, duckweeds may then be less hormonally disrupted and grow less.

One pervasive theme across studies of leachate from tire wear particles, is that not all tires, organisms, and methods are equivalent. Diverse methods find diverse biological effects ranging from acute lethality in coho salmon (Peter et al., 2018) to no effects at even high doses for some invertebrates (Redondo-Hasselerharm et al., 2018). Studies on tire particles vary in a number of factors that may influence toxicity, not exhuastively including: the brand of tire (Wik and Dave, 2006), size of tire particles used (though perhaps not always, Rhodes et al., 2012; Khanal et al., 2014), the age of the tire (Day et al., 1993; Sharma et al., 2010), whether or not road wear is simultaneously considered (Redondo-Hasselerharm et al., 2018), whether or not the particles or leachate or both are tested (Khan et al., 2019), leaching time (Wik and Dave, 2006; Rhodes et al., 2012), leaching conditions (Marwood et al., 2011) and leachate storage (Khanal et al., 2019). It is hard to predict how differences in methods, tires, or study species might have affected our results, but we note that the studies on similar organisms above do not find opposing signs of effects across factors, but rather stronger and weaker effects (for example, Panko et al., 2013; Kolomijeca et al., 2020). Finally, we note commonalities in the effects of temperature. Kolomijeca et al. (2020) similarly observed stronger effects of leachate from tire wear particles at high temperature on fathead minnow, and Marwood et al. (2011) observed that leachate produced at higher temperatures had greater effects, though these studies both observed stronger negative effects, while we observed stronger positive effects of temperature and tire when duckweed were inoculated with microbiomes. Broadening our view, it appears that temperature exacerbation of contaminant effects may be a fairly common outcome across contaminants in aquatic systems (Crain et al., 2008; Jackson et al., 2016).

## Conclusions

Changing environments can disrupt species interactions, and duckweed-microbiome interactions are no exception: microbiomes were more costly to duckweed with tire particle leachate or warming, and leachate may shift selection pressures towards microbes that reduce duckweed growth (or duckweeds that reduce microbial growth). Further, with our finding that some effects of leachate from tire particles depended on temperature or CO_2_, our study joins a vast array of cases where multiple global change factors have non-additive effects (Darling and Côté, 2008; Crain et al., 2008; Jackson et al., 2016). With an ever increasing suite of anthropogenic stressors comes more potential for such “ecological surprises,” including for disrupted species interactions. Effects of anthropogenic contaminants on species interactions therefore continues to be a critical area of research: shifted strengths or flipped signs of interaction outcomes could echo through food webs and alter basic ecosystem functions from productivity to stability.

## Author contributions

All authors contributed substantially to the design of the study, provisioning of materials, and/or revising of the manuscript.

## Acknowledgements

The authors would like to thank students and volunteers who have contributed to maintaining duckweed and microbe cultures in the lab, as well as members of the Frederickson, Rochman, and Sinton labs for helpful discussions.

This work was primarily supported through the Strategic Projects Grant Program of the Natural Science and Engineering Research Council of Canada (NSERC) (to DS and CMR, STPGP 506882). Other funding included NSERC Discovery Grants (to DS, CMR, and MEF), an Alexander Graham Bell Canada Graduate Scholarship-Doctoral from the NSERC (TFL), an E.W.R. Steacie Memorial Fellowship (DS), and the Canada Research Chairs Program (DS). Some of the equipment used in this study was supported by the 3D (Diet, Digestive Tract, and Disease) Centre funded by the Canadian Foundation for Innovation and Ontario Research Fund, project numbers 19442 and 30961.

## Data Accessibility

Data will be accessible upon publication via figshare and Github. Scripts will also be accessible on Github.

---

**Figure S1:**
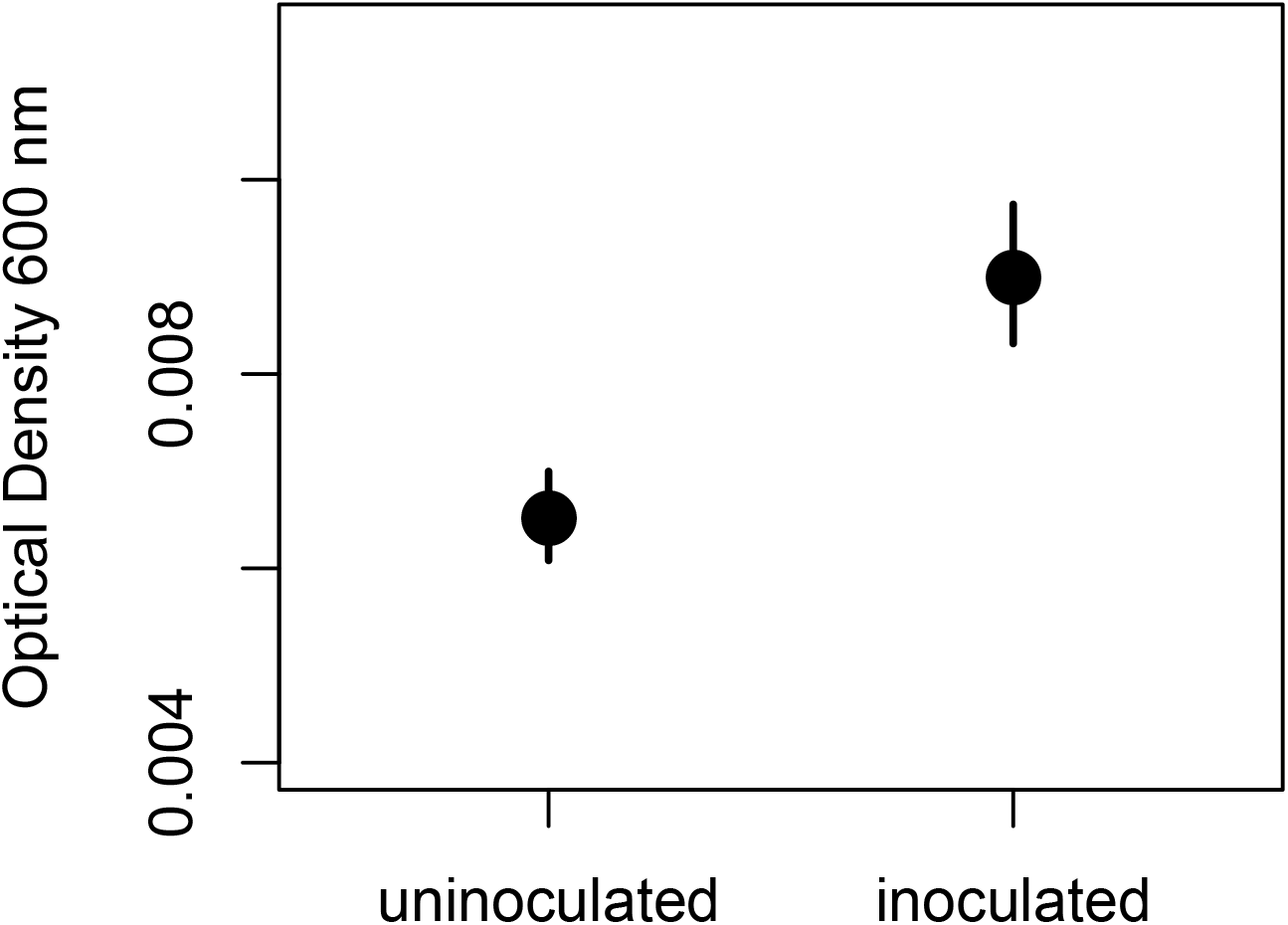
Effect of inoculation with microbiome on total microbial growth (optical density). Means (points) and standard errors (bars) are shown.

**Figure S2:**
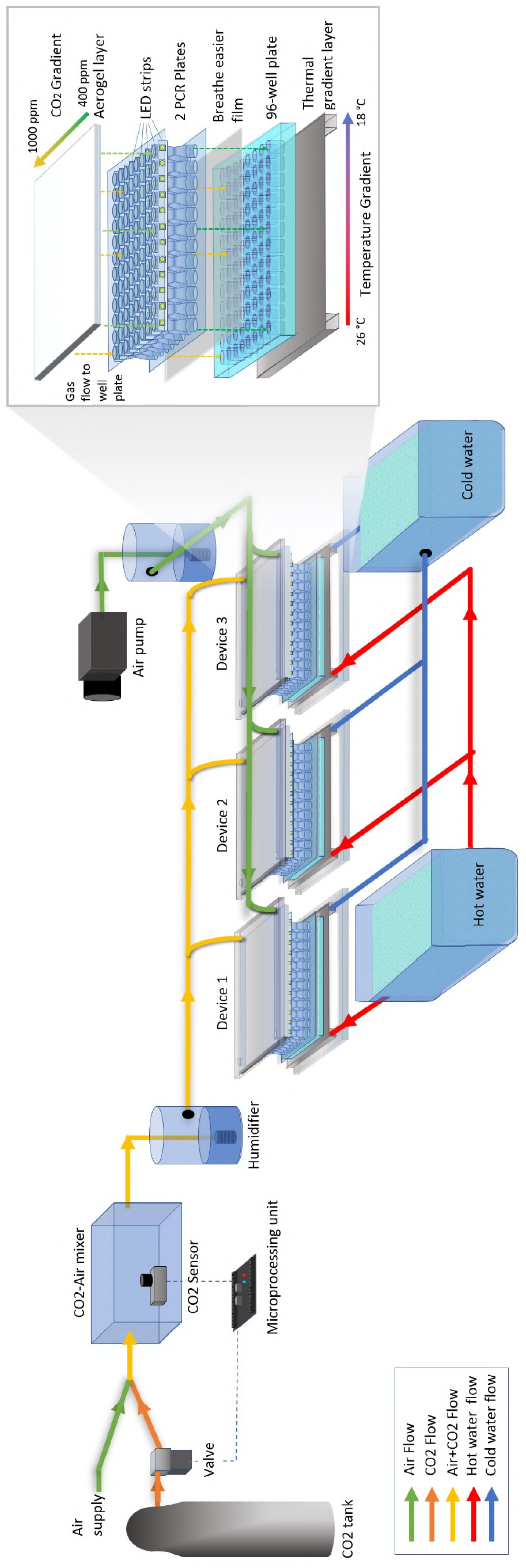
Graphical representation of device setup. Briefly, a target elevated CO_2_ concentration for the highest treatment (yellow) was achieved by mixing pure CO_2_ (orange) and air (green) via the controlling action of a microprocessing unit, valve and sensor (see text and Nguyen et al., 2018; Yang et al., 2020). Then, both this mix and air (green) were humidified and delivered to the plate on opposite sides, and an even gradient was generated with an aerogel. Below this aerogel layer, we placed tubes that connect the aerogel gas gradient to experimental wells of a 96-well plate. LED light strips were interlaced in the spacing between these tubes. Next, we placed a breathable, cell-barrier membrane over the wells. Finally, below the plate, we applied a temperature gradient using an aluminum plate with hot and cold water piped across either end (orthogonal to the gas gradient, see inset, Methods text). The three plates in parallel depicted here show one replicate, which was repeated three times. The schematics of one of the three plates is shown in inset on the right, with the different layers shown. The three leachate treatments would be randomly assigned across plates within a replicate and are therefore not shown.

**Figure S3:**
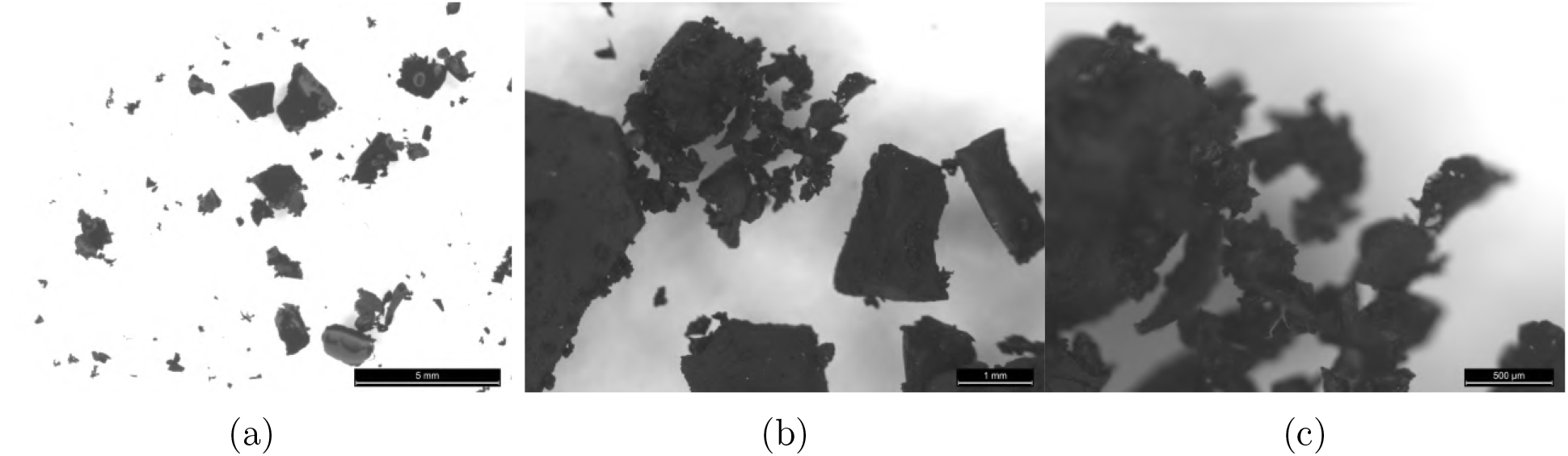
Characteristic images of tire particles generated and used in the experiment. a) tire particles in surfactant, removing most static attractions and images blown out as described in Methods. b) tire particles without surfactant. c) a region of the same particles at higher magnification.

**Figure S4:**
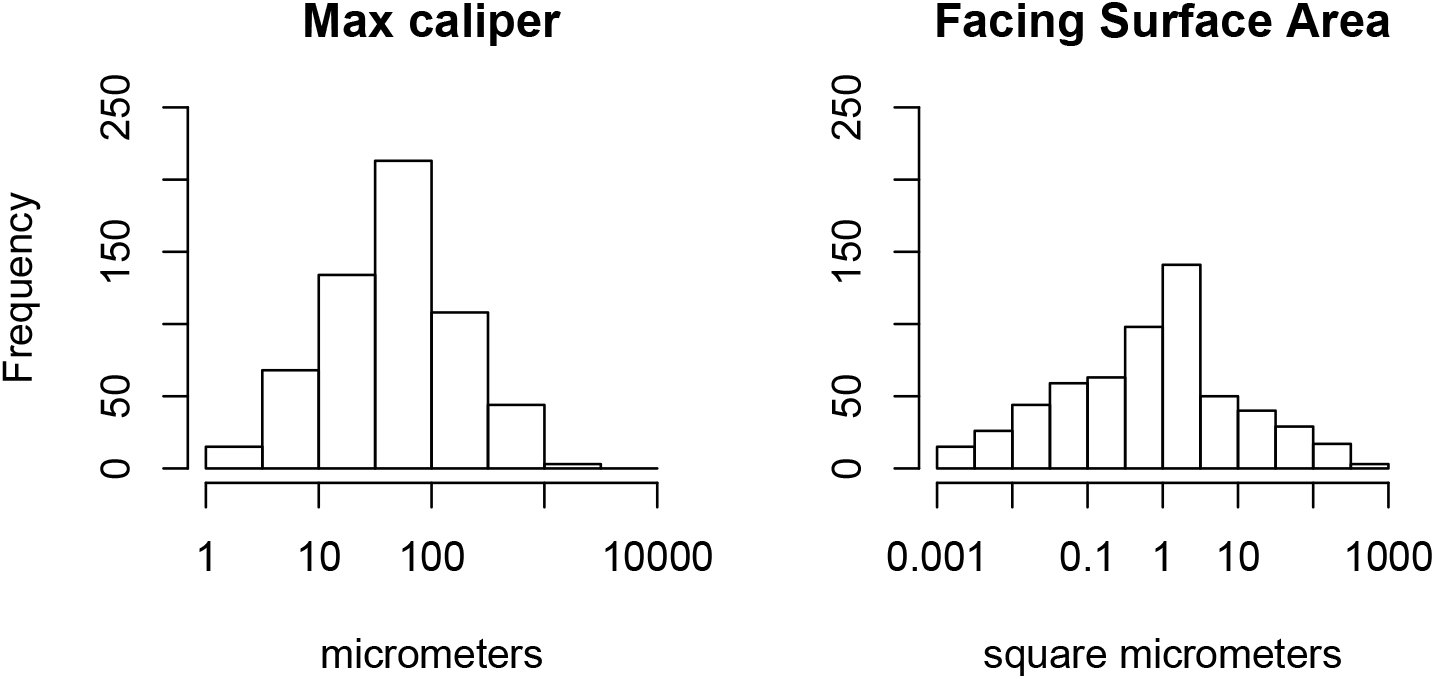
Distribution of tire particle size measurements. The maximum caliper (Feret’s Diameter, left) and the top surface area (visible in the in the image, right).

**Figure S5:**
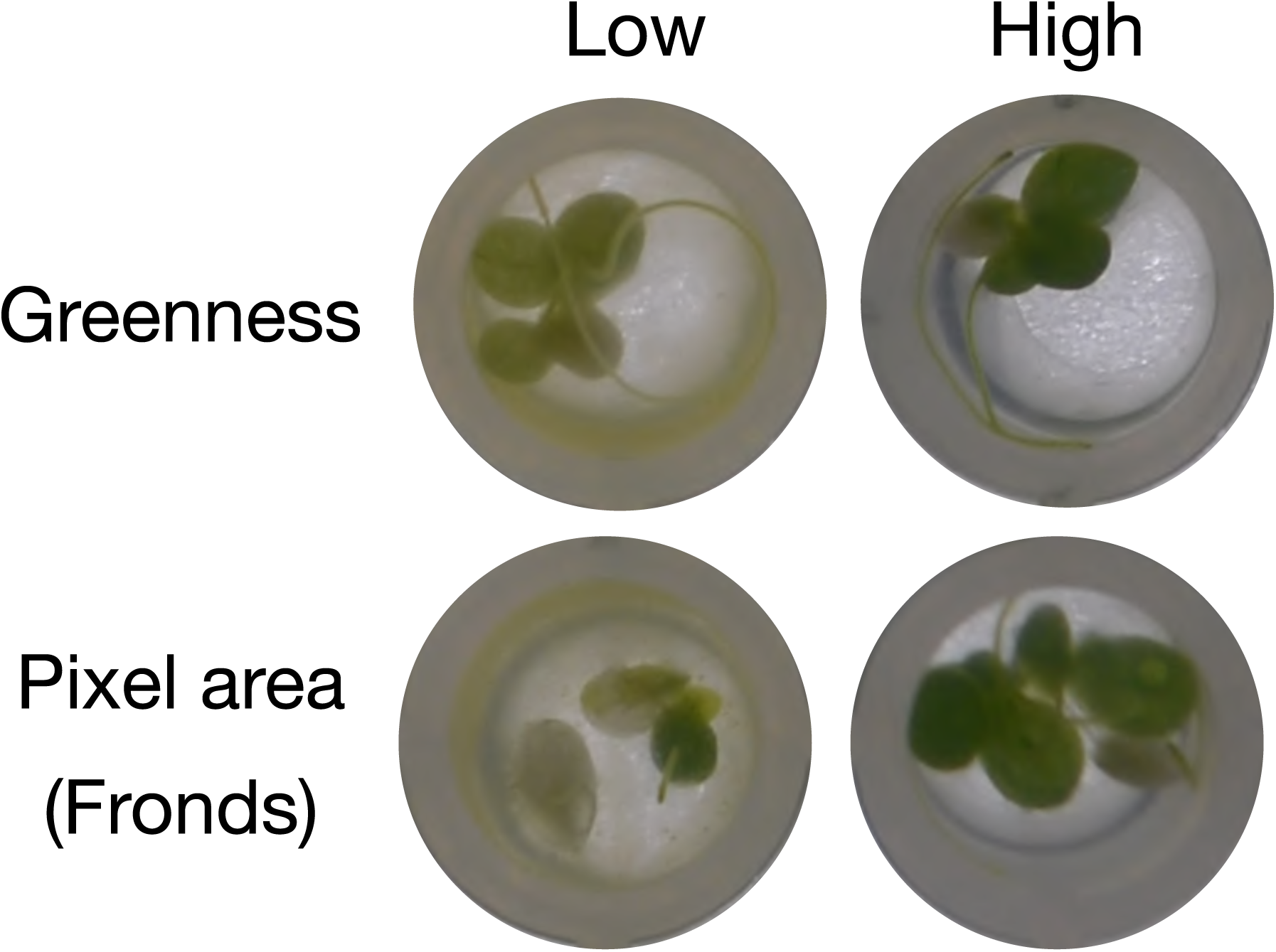
Example images of duckweed from the experiment, for duckweeds scored in ImageJ to have lower and higher greenness and pixel area (related to frond number). These are from the third round, and are of the plate from the tire particle leachate treatment of 10 g/L concentration.

